# Integration of Alzheimer’s disease genetics and myeloid genomics reveals novel disease risk mechanisms

**DOI:** 10.1101/694281

**Authors:** Gloriia Novikova, Manav Kapoor, Julia TCW, Edsel M. Abud, Anastasia G. Efthymiou, Haoxiang Cheng, John F. Fullard, Jaroslav Bendl, Panos Roussos, Wayne W. Poon, Ke Hao, Edoardo Marcora, Alison M. Goate

## Abstract

Genome-wide association studies (GWAS) have identified more than thirty loci associated with Alzheimer’s disease (AD), but the causal variants, regulatory elements, genes and pathways remain largely unknown thus impeding a mechanistic understanding of AD pathogenesis. Previously, we showed that AD risk alleles are enriched in myeloid-specific epigenomic annotations. Here, we show that they are specifically enriched in active enhancers of monocytes, macrophages and microglia. We integrated AD GWAS signals with myeloid epigenomic and transcriptomic datasets using novel analytical approaches to link myeloid enhancer activity to target gene expression regulation and AD risk modification. We nominate candidate AD risk enhancers and identify their target causal genes (including AP4E1, AP4M1, APBB3, BIN1, CD2AP, MS4A4A, MS4A6A, PILRA, RABEP1, SPI1, SPPL2A, TP53INP1, ZKSCAN1, and ZYX) in sixteen loci. Fine-mapping of these enhancers nominates candidate functional variants that likely modify disease susceptibility by regulating causal gene expression in myeloid cells. In the MS4A locus we identified a single candidate functional variant and validated it experimentally in human induced pluripotent stem cell (hiPSC)-derived microglia. Combined, these results strongly implicate dysfunction of the myeloid endolysosomal system in the etiology of AD.

## Introduction

Alzheimer’s disease (AD) is the most common type of dementia with a global burden of approximately 50 million people and no disease-modifying treatments available ^1^. Several lines of genetic evidence implicate myeloid cells in the etiology of AD ^2^. Whole-exome sequencing and microarray studies have identified rare coding variants associated with AD in genes (e.g., TREM2 ^3^, SORL1 ^4^, ABI3 ^5^, PLCG2 ^5^ and ABCA7 ^6^) that play important roles in myeloid cells of the brain (microglia) and peripheral tissues (e.g., monocytes and macrophages) and have high relative expression levels in microglia compared to other brain cell types ^7^. Genome-wide association studies (GWAS) have identified common non-coding variants associated with AD in more than thirty loci, but the identification of causal variants and genes in these loci is still lacking. Earlier studies have focused on mapping genes to AD risk loci using whole-blood and brain expression quantitative trait loci (eQTL) datasets ^8–10^. However, using tissue-level data poses obstacles to identifying myeloid-specific signals, because myeloid cells (microglia and monocytes) represent small fractions (∼10%) of the total cell population in their respective tissues (brain and peripheral blood). More importantly, given the strong enrichment of AD risk alleles in myeloid-specific epigenomic annotations and expressed genes ^11, 12^, it is imperative to investigate the role of myeloid epigenomes and transcriptomes in the modulation of AD susceptibility.

In this study, we investigated the effects of AD risk variants on the epigenome and transcriptome of myeloid cells. We first show that AD risk alleles are specifically enriched in active enhancers in monocytes, macrophages and microglia and identify transcription factor binding motifs (TFBMs) overrepresented within these regulatory elements. We further identify myeloid transcription factors (TFs) whose binding sites at active enhancers are burdened by AD risk variants. Given the selective enrichment of AD risk alleles in myeloid active enhancers, we sought to link the activity of myeloid enhancers that contain AD risk variants to target gene expression regulation and AD risk modification. To accomplish this we used two complementary approaches.

First, we mapped myeloid active enhancers that contain AD risk alleles to their target genes by integrating chromatin interactions (promoter-capture Hi-C) and eQTL datasets from monocytes and macrophages. This approach allowed us to nominate candidate causal genes in twelve genome-wide significant and two suggestive AD risk loci, TP53INP1 and APBB3. In our second approach, we used Summary data-based Mendelian Randomization (SMR) ^13^ to investigate the causal relationship between enhancer activity, target gene expression regulation and AD risk modification. This approach allowed us to identify specific myeloid active enhancers that likely modify AD risk by regulating the expression of their target genes in ten loci. Importantly, the target genes of the myeloid active enhancers identified by these two analytical approaches were highly consistent and implicate the endolysosomal system of myeloid cells in the etiology of AD. We further fine-mapped these AD risk enhancers to identify candidate functional variants that affect TF binding, modulate enhancer activity and regulate causal gene expression in eight loci, and experimentally validated one of these candidate causal variants in the MS4A locus in human induced pluripotent stem cell (hiPSC)-derived microglia.

## Results

### AD risk alleles are specifically enriched in active enhancers of monocytes, macrophages and microglia

Our earlier analyses showed a significant enrichment of AD risk alleles in various myeloid-specific epigenomic annotations, but not in brain or other tissues ^11^. To further dissect this enrichment, we used ChIP-Seq profiles of histone modifications that define the chromatin signatures of regulatory elements (H3K27ac for active enhancers and promoters, H3K4me1 for enhancers, and H3K4me2 for active enhancers and promoters) from monocytes, macrophages and microglia to annotate the genome with myeloid active enhancers (AE), active promoters (AP), primed enhancers (PE) and primed promoters (PP) (see Methods) ^14^. To identify which of these myeloid regulatory elements are enriched for AD risk alleles, we performed stratified LD score regression (LDSC) ^15^ of AD single nucleotide polymorphism (SNP) heritability partitioned by the aforementioned epigenomic annotations using the International Genomics of Alzheimer’s Project (IGAP) AD GWAS dataset ^16^. This analysis revealed selective enrichment of AD risk alleles in active enhancers of monocytes, macrophages and microglia (Figure 1a). In contrast, schizophrenia SNP heritability (using the Psychiatric Genomics Consortium SCZ GWAS dataset as control ^17^) was not enriched in any of these myeloid regulatory elements (Figure 1a).

**Figure 1.**
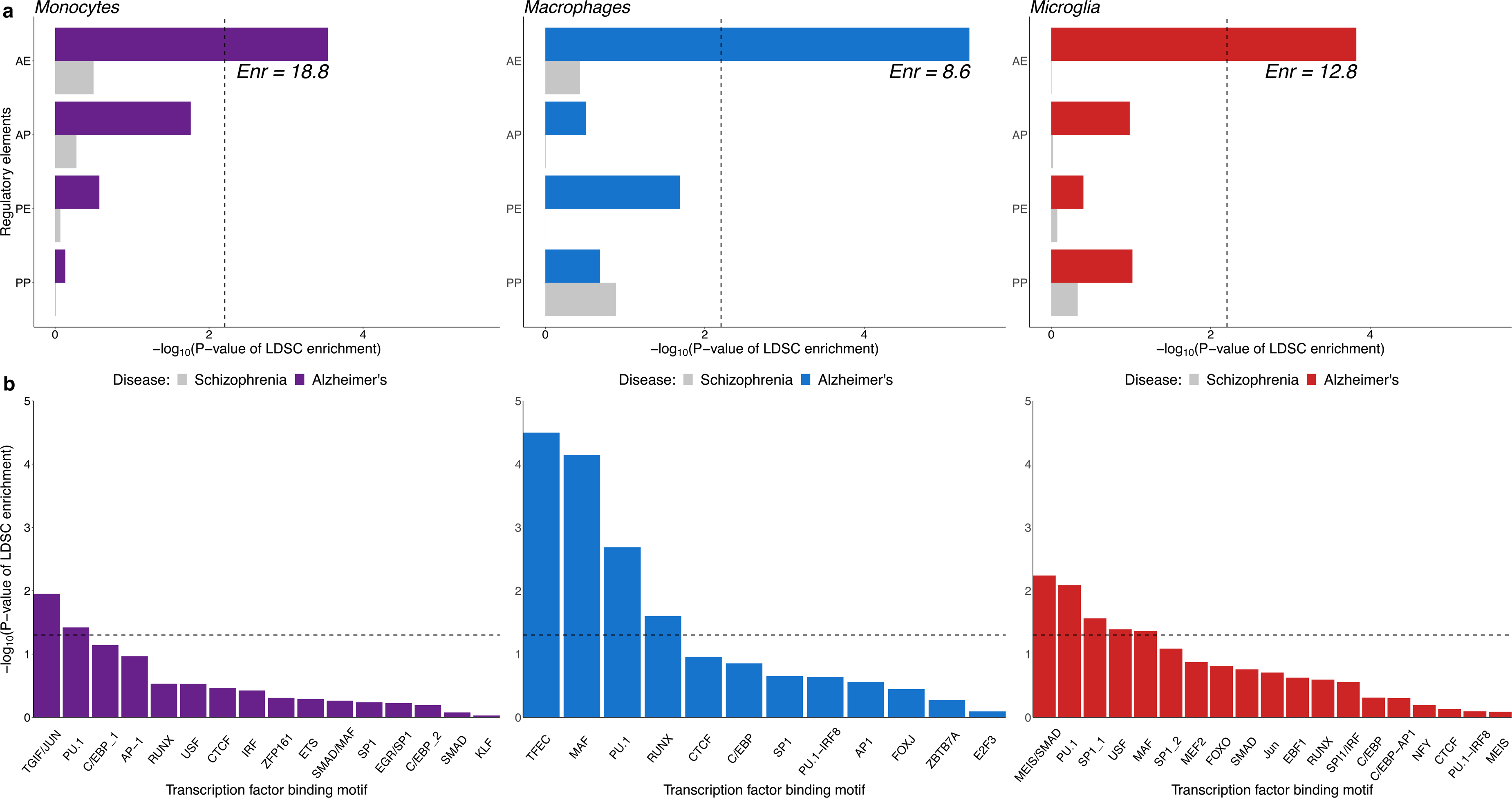
AD risk alleles are specifically enriched in myeloid active enhancers and in putative transcription factor binding sites located in these enhancers. a. -Log10 of enrichment P-values obtained from stratified LD Score Regression (LDSC) analysis of AD SNP heritability partitioned by active enhancer (AE), active promoter (AP), primed enhancer (PE) and primed promoter (PP) annotations in monocytes, macrophages and microglia. Enr = Enrichment of AD SNP heritability partitioned by active enhancer annotations. Dashed line indicates Bonferroni-corrected significance threshold. The enrichment standard errors are 4.8, 1.7 and 3.2 for monocytes, macrophages and microglia, respectively. b. -Log10 of enrichment P-values obtained from stratified LD Score Regression (LDSC) analysis of AD SNP heritability partitioned by ATAC-Seq subsets. The subsets were obtained by stratifying ATAC-Seq regions in monocytes, macrophages and microglia by the presence of the binding motif of TFs (listed on the x-axis) that were found to be overrepresented in active myeloid enhancers and expressed in microglia (TPM≥1)^14^.

To identify TFs that likely regulate the activity of myeloid enhancers, we performed de novo motif analysis ^18^ in open chromatin regions (identified by ATAC-Seq) that overlap with active enhancers in all three cell types (Supplementary Table 1). The binding motif for PU.1 (a transcription factor critical for myeloid and B-lymphoid cell development and function and an AD risk gene (SPI1) ^11^) was the best match for the most highly overrepresented sequence motif in active enhancers across all three cell types, followed by C/EBP, CTCF and RUNX binding motifs. The binding motif for MEF2 family TFs (which includes MEF2C in another AD risk locus ^16^) was the best match for the most highly overrepresented sequence motif in active enhancers of human microglia, consistent with findings in mouse microglia ^19^. To test whether the binding sites of TFs that likely regulate active myeloid enhancers are enriched for AD risk variants, we stratified ATAC-Seq regions in all three cell types by the presence of the binding motifs of the TFs that were found to be overrepresented in active myeloid enhancers, and applied LDSC to quantify the enrichment of AD SNP heritability partitioned by these subsets of ATAC-Seq regions (Figure 1b). ATAC-Seq regions overlapping with active enhancers that were positive for the PU.1 binding motif in all three cell types were enriched for AD risk alleles. MAF binding motif-positive ATAC-Seq regions were enriched for AD risk alleles in macrophage and microglial active enhancers. SMAD, USF and SP1 binding motif-positive ATAC-Seq regions were enriched for AD risk alleles only in microglial active enhancers. Interestingly, a study comparing two mouse strains reported that genetic variants in Mafb, Smad3, and Usf1 binding sites affected PU.1 binding specifically in microglia, suggesting that these TFs could be binding partners of PU.1 in microglia ^20^. These results show that AD risk alleles are specifically enriched in active enhancers of monocytes, macrophages and microglia, and nominate shared and cell-type specific TFs that likely regulate the activity of these regulatory elements. Additionally, these results implicate TFs whose binding to myeloid active enhancers is likely to be affected by AD risk alleles. These results support our hypothesis that TF binding sites might be altered by AD risk variants to affect myeloid enhancer activity and gene expression, which in turn modulate disease susceptibility by altering the biology of myeloid cells.

### Integration of AD GWAS signals with myeloid epigenomic annotations, chromatin interactions (promoter-capture Hi-C) and eQTL datasets identifies candidate causal genes in fourteen AD risk loci

Promoter-enhancer interactions constitute one of the most fundamental mechanisms of gene expression regulation, where enhancer elements are brought into close proximity to cognate promoters to stimulate transcription of their target genes ^18^. Given the observed enrichment of AD risk alleles in myeloid active enhancers, we reasoned that harnessing information about the spatial organization of chromatin and integrating it with epigenomic annotations and eQTLs in myeloid cells would facilitate the identification of candidate causal genes regulated by these elements in AD risk loci. As chromatin interactions and eQTL datasets are currently not available for human microglia and our partitioned AD SNP heritability estimates suggest that peripheral myeloid cells are good proxy cell types for microglia in the brain, we used datasets from human peripheral blood monocytes and monocyte-derived macrophages as we did previously ^11^. We first identified active enhancers in monocytes and macrophages that contain AD risk alleles (P≤1×10^-6^). We then selected active enhancers that interact with at least one promoter of genes expressed in microglia (TPM ≥1) ^14^ and contain AD risk variants that are eQTLs for the same gene in monocytes and macrophages (FDR≤5%) using the Javierre et al. 2016 promoter-capture Hi-C dataset ^18^ and the Cardiogenics ^21^ and Fairfax ^22^ eQTL datasets, hereafter referred to as AD risk enhancers. Using this approach we nominate candidate causal genes in fourteen genome-wide significant and suggestive AD risk loci (Table 1). In some loci, this analysis identified genes that have known AD-associated coding variants (ABCA7 ^23^) and genes that have been identified as most likely causal in previous studies (BIN1 ^24^ and PTK2B ^25^). In other loci, we uncovered co-regulation of the expression of multiple target genes by shared AD risk enhancers. For example, in the SPI1 locus, we identified AD risk enhancers shared by ACP2, MADD, MYBPC3, NR1H3, NUP160 and SPI1 in monocytes and macrophages. Similarly, in the ZCWPW1 locus, we identified AD risk enhancers shared by AP4M1, PILRA, PILRB, and ZCWPW1 in monocytes, and by AP4M1, MCM7, PILRA, PILRB, PVRIG, STAG3 and ZCWPW1 in macrophages. This could reflect either multiple causal genes within these loci or a single causal gene and several risk neutral genes that show association by virtue of expression co-regulation. Additional evidence is necessary to distinguish between these two possibilities and prioritize one or more genes in the locus as we have shown for SPI1 at the respective (previously CELF1) locus ^11^.

**Table 1.** Candidate causal genes identified through integration of AD GWAS signals with myeloid active enhancer annotations, promoter-capture Hi-C, and eQTLs datasets.

| **Locus** | **Monocytes** | **Monocyte-derived macrophages** |
| --- | --- | --- |
| *BIN1* | *BIN1* | *BIN1* |
| *SPI1* (previously *CELF1*) | *ACP2, FNBP4, MADD, MYBPC3, MTCH2, NR1H3, NUP160, PSMC3, SPI1* | *MTCH2, MYBPC3, NUP160, PSMC3, SPI1* |
| *ZYX* (previously *EPHA1*) | *ZYX* | *ZYX* |
| *MS4A* | *MS4A6A* | *MS4A6A* |
| *PILRA* (previously *ZCWPW1*) | *AP4M1, GATS, PILRA, PILRB, ZCWPW1* | *AP4M1, GATS, MCM7, MOSPD3, PILRA, PILRB, PVRIG, STAG3, TRIM4, ZCWPW1* |
| *TP53INP1* (previously *NDUFAF6*) | *INTS8, TP53INP1* | *INTS8, TP53INP1* |
| *AP4E1/SPPL2A* | - | *AP4E1* |
| *RIN3* (previously *SLC24A4* locus) | *RIN3* | - |
| *ABCA7* | *ABCA7, CNN2, CIRBP* | *ABCA7, CNN2, GPX4, WDR18* |
| *APBB3* (previously *HBEGF* locus) | - | *APBB3, PFDN1* |
| *RABEP1* (previously *SCIMP* locus) | - | *CHRNE* |
| *PTK2B* | *PTK2B* | - |
| *CASS4* | *AURKA* | - |
| *TREM2* | *NFYA* | - |

Additionally, these analyses revealed regulatory landscapes that are shared across cell types or are cell type-specific. In the BIN1 locus, we observed conserved AD risk enhancer-promoter chromatin interactions and similar eQTL signal profiles in monocytes and macrophages, suggesting that the AD risk regulome is similar in these two cell types and points to BIN1 as the strongest candidate causal gene at this locus (Figure 2a). Conversely, in the ZYX (previously EPHA1) locus, we observed stronger chromatin interactions with a ZYX promoter in macrophages (mean interaction score 3.9 and 8.5 in monocytes and macrophages, respectively) and different eQTL signal profiles between monocytes and macrophages, suggesting that the AD risk regulome is different in these two cell types albeit pointing to the same candidate causal gene (Supplementary Figure 1). Finally, we identified candidate causal genes, TP53INP1 (Figure 2b) and APBB3 in suggestive loci (P≤1×10^-6^). In summary, this approach allowed us to nominate candidate causal genes in twelve genome-wide significant and two suggestive AD risk loci.

**Figure 2.**
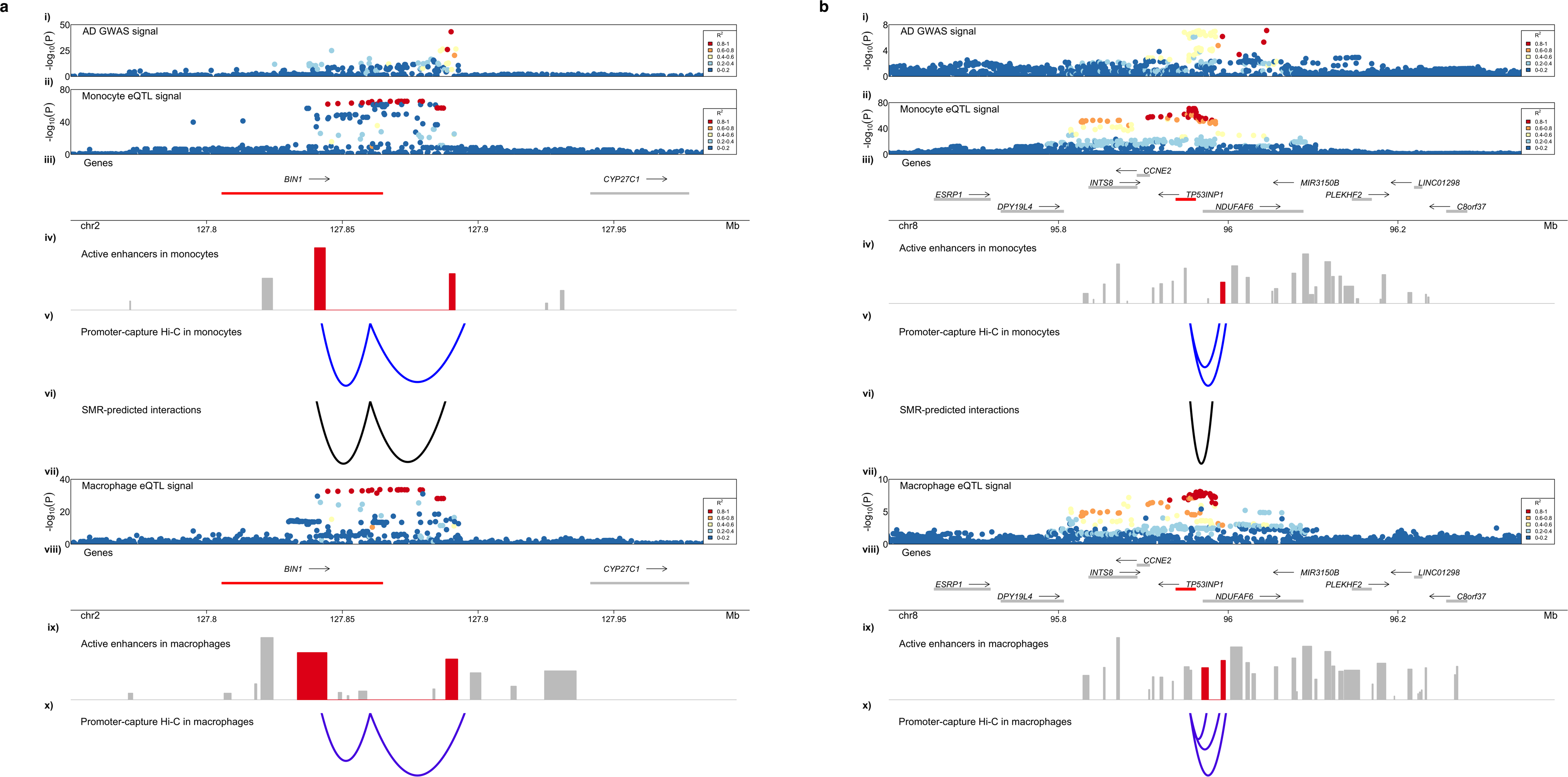
AD risk enhancers interact with the promoters of BIN1 and TP53INP1. **a.** i) AD GWAS signal in the BIN1 locus. ii) eQTL signal for BIN1 in monocytes obtained from the Cardiogenics study. iii) Genes that reside in the locus are plotted. Putative AD risk genes are highlighted in red. The arrow indicates the direction of transcription, while the bar indicates the gene body. iv) Active enhancers in monocytes are plotted. Putative AD risk enhancers are highlighted in red. v) Promoter-capture Hi-C interactions between the BIN1 promoter and AD risk enhancers in monocytes. vi) Enhancer-gene interactions predicted by SMR analysis of causal associations between enhancer activity and BIN1 expression in monocytes. vii) eQTL signal for BIN1 in macrophages obtained from the Cardiogenics study. viii) Genes that reside in the locus are plotted. Putative AD risk genes are highlighted in red. The arrow indicates the direction of transcription, while the bar indicates the gene body. ix) Active enhancer elements in macrophages are plotted. Putative AD risk enhancers are highlighted in red. x) Promoter-capture Hi-C interactions between the BIN1 promoter and AD risk enhancers in macrophages. b. i) AD GWAS association signal in the TP53INP1 locus. ii) eQTL signal for TP53INP1 in monocytes obtained from the Cardiogenics study. iii) Genes that reside in the locus are plotted. Putative AD risk genes are highlighted in red. The arrow indicates the direction of transcription, while the bar indicates the gene body. iv) Active enhancers in monocytes are plotted. Putative AD risk enhancers are highlighted in red. v) Promoter-capture Hi-C interactions between the TP53INP1 promoter and AD risk enhancers in monocytes. vi) Enhancer-gene interactions predicted by SMR analysis of causal associations between enhancer activity and TP53INP1 expression in monocytes. vii) eQTL signal for TP53INP1 in macrophages obtained from the Cardiogenics study. viii) Genes that reside in the locus are plotted. Putative AD risk genes are highlighted in red. The arrow indicates the direction of transcription, while the bar indicates the gene body. ix) Active enhancer in macrophages are plotted. Putative AD risk enhancers are highlighted in red. x) Promoter-capture Hi-C interactions between the TP53INP1 promoter and AD risk enhancers in macrophages.

### Integration of AD GWAS signals with myeloid epigenomic annotations, chromatin activity (hQTL) and eQTL datasets identifies candidate causal genes in ten AD risk loci

Although chromatin interactions between active enhancers and gene promoters may suggest target gene expression regulation, inferring causal relationships between chromatin activity at enhancer elements and target gene expression can provide additional evidence for such regulation and help identify genetic variants that mediate these relationships to modulate disease susceptibility. We used SMR to explore the causal path that links myeloid enhancer activity to target gene expression regulation and AD risk modification. To accomplish this, we used datasets from monocytes ^26^, since chromatin activity QTLs (hQTLs) are currently not available for human microglia or other macrophages. We first identified active enhancers in monocytes that contain AD risk alleles (P≤1×10^-6^) and hQTLs (genetic variants affecting the activity of the active enhancer at FDR≤10%) and used coloc ^27^ to select those with evidence of independent or colocalized AD GWAS and hQTL signals (PP.H3.abf + PP.H4.abf≥0.5) (Supplementary Table 2). To investigate the link between myeloid enhancer activity and target gene expression regulation, we used SMR to test for causal association between hQTL and eQTL effects in monocytes at the 24 active enhancers selected using coloc. We identified multiple genes that are likely regulated by these AD risk enhancers (Figure 3a, Table 2, Supplementary Table 3), including BIN1, CD2AP, GPR141, MS4A4A, MS4A6A, RABEP1, SPI1, TP53INP1 and ZYX. We then used SMR to test for causal association between the expression of genes regulated by the AD risk enhancers identified above and disease susceptibility. These analyses revealed specific enhancers in monocytes, whose activity is causally associated with expression of their target genes, which in turn is causally associated with AD risk, including BIN1, GPR141, MS4A4A, MS4A6A, RABEP1, SPI1, TP53INP1 and ZYX (Figure 3b, Supplementary Table 4). Twelve of twenty-two genes nominated through causal associations between chromatin activity and gene expression and eight of thirteen genes nominated through causal associations between gene expression and disease susceptibility identified using the Cardiogenics monocyte eQTL dataset were replicated using the Fairfax monocyte eQTL dataset (Supplementary Tables 5-6). Since the replication cohort is smaller, we expect that a larger number of associations would replicate in a larger cohort, given the fact that all genes found through associations using the Fairfax dataset were significant in the main analysis using the Cardiogenics dataset. Additionally, the AD risk enhancers for BIN1, SPI1, TP53INP1 and ZYX identified as described above by SMR also interact with the promoters of the same genes, providing converging evidence for causal target gene expression regulation by these AD risk enhancers (Figure 2a, iv-vi; Figure 2b, iv-vi).

**Figure 3.**
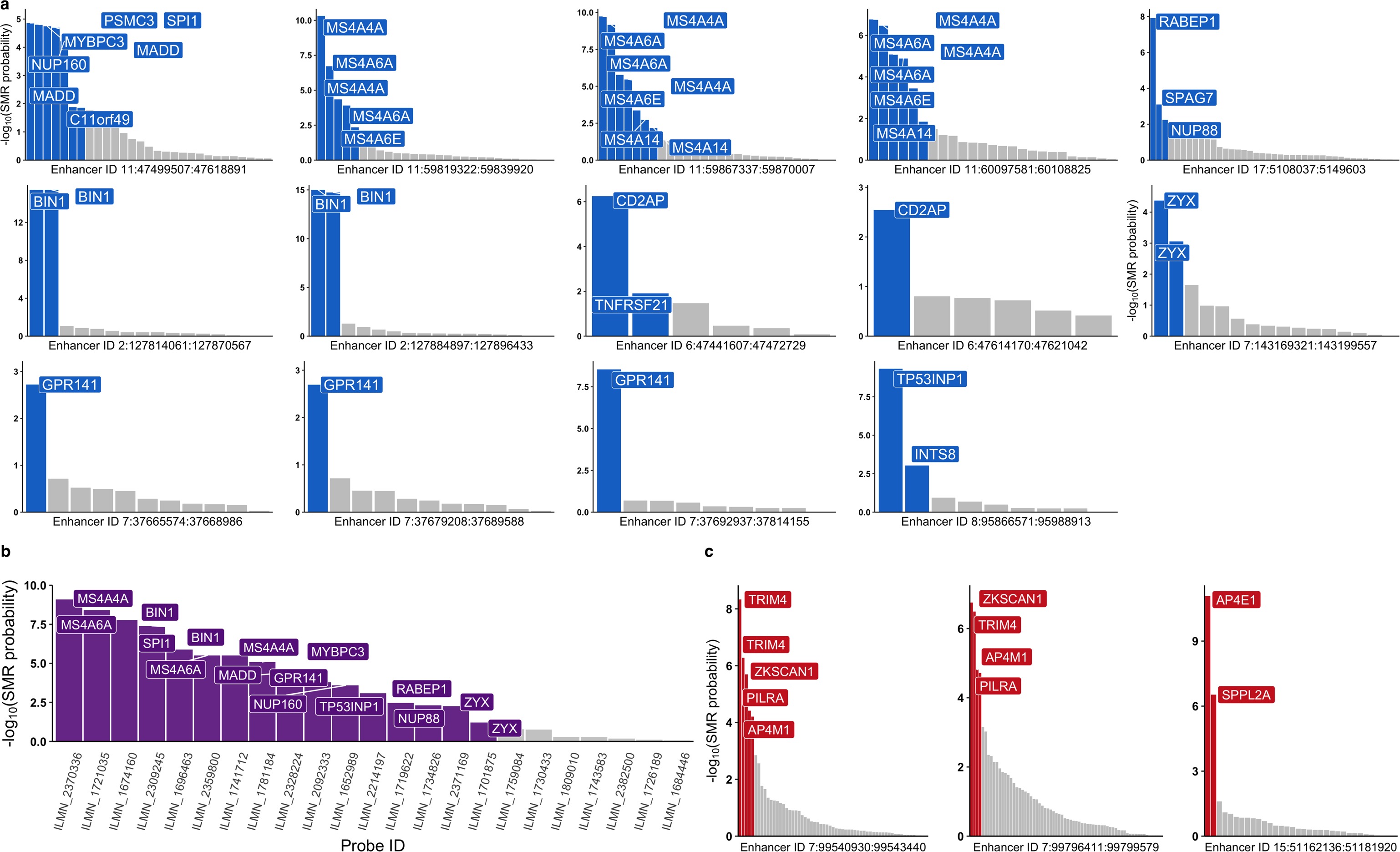
Causal associations between myeloid enhancer activity, target gene expression regulation and AD risk modification point to candidate causal genes. a. -Log10 of causal association probabilities between enhancer activity and gene expression in monocytes obtained through SMR analysis for each probe are plotted for each active enhancer element. Probes (labeled by the respective gene) in blue indicate significant associations, while grey bars indicate non-significant associations based on a 10% FDR threshold. b. -Log10 of causal association probabilities between gene expression and AD risk. Probes (labeled by their respective gene) in purple indicate significant associations, while grey bars indicate non-significant associations based on a 10% FDR threshold. c. -Log10 of causal association probabilities between activity of two enhancers in the PILRA locus and one enhancer in the SPPL2A locus and gene expression in monocytes obtained through SMR analysis for each probe are plotted. Probes (labeled by the respective gene) in red indicate significant associations, while grey bars indicate non-significant associations based on a 10% FDR threshold.

**Table 2.** Candidate causal genes identified through integration of AD GWAS signals with myeloid enhancer annotations, hQTL, and eQTL datasets.

| **Locus** | **Genes implicated through enhancer to gene associations** | **Genes implicated through enhancer to gene expression to disease risk associations** |
| --- | --- | --- |
| *BIN1* | *BIN1* | *BIN1* |
| *SPI1* | *C11orf49, MADD, MYBPC3, NUP160, PSMC3, SPI1* | *MADD, MYBPC3, NUP160, SPI1* |
| *CD2AP* | *CD2AP* | - |
| *ZYX* (previously *EPHA1*) | *ZYX* | *ZYX* |
| *GPR141* (previously *NME8*) | *GPR141* | *GPR141* |
| *TP53INP1* | *INTS8, TP53INP1* | *TP53INP1* |
| *MS4A* | *MS4A14, MS4A4A, MS4A6A, MS4A6E\** | *MS4A4A, MS4A6A* |
| *RABEP1* (previously *SCIMP*) | *NUP88, RABEP1, SPAG7* | *NUP88, RABEP1* |
| *PILRA* <sup>a</sup> (previously *ZCWPW1*) | *AP4M1, TRIM4, ZKSCAN1, PILRA* | *AP4M1, ZKSCAN1, PILRA* |
| *AP4E1/SPPL2A* <sup>b</sup> | *AP4E1, SPPL2A* | *SPPL2A* |

<sup>a</sup>Association reported with primed enhancer
<sup>b</sup>Association reported with a distal active enhancer that does not contain AD risk variants, but has hQTLs that colocalize with AD risk alleles
\*Not expressed in microglia

Although we observed a global enrichment of AD risk alleles in myeloid active enhancers across the human genome (Figure 1a), we discovered a small subset of loci where the regulatory elements associated with causal gene expression regulation are not active enhancers. For example, we identified two primed enhancers in monocytes whose hQTLs are causally associated with expression of PILRA, AP4M1 and ZKSCAN1, which is in turn causally associated with AD risk (Figure 3c). Moreover, we identified an active enhancer element whose activity is regulated by AD risk alleles located at a distance from it and which is strongly associated with expression of AP4E1 and SPPL2A in monocytes (Figure 3c). In turn, expression of SPPL2A is causally associated with AD risk (Figure 3c). Furthermore, this active enhancer interacts with the promoter of SPPL2A, providing converging evidence for causal regulation of SPPL2A expression by this enhancer element. Therefore, it is possible that AD risk alleles indirectly affect the activity of this enhancer by functional coupling through chromatin looping or another mechanism.

### Fine-mapping using myeloid epigenomic annotations identifies candidate causal variants in eight AD risk loci

To prioritize candidate causal variants in AD risk enhancers we selected loci where we discovered significant associations between enhancer activity, gene expression and AD risk (i.e. BIN1, GPR141, MS4A, PILRA/AP4M1, RABEP1, SPI1, SPPL2A/AP4E1, TP53INP1, and ZYX). We first selected variants in high to moderate LD (R^2^=0.8) with the tagging variant in each locus and queried them in Haploreg ^28^ to identify coding variants. We identified a missense variant (rs1859788-A) in PILRA that is in high LD with the tagging variant (R^2^=0.85) and was previously shown to alter the ligand binding affinity of PILRA ^29^. Conditioning on this variant eliminates the AD GWAS signal at this locus (Supplementary Figure 2). The other eight AD risk loci did not contain coding variants in high LD with the tagging variant, prompting us to proceed with fine-mapping to prioritize candidate non-coding functional variants. To accomplish this we used PAINTOR, a Bayesian fine-mapping method that allows for integration of epigenomic annotations ^30^. Due to the inflation of posterior probabilities when GWAS and individual-level genotype data are not matched ^31^, we used summary-level GWAS statistics and matched individual-level genotype data from the <u>Alzheimer’s Disease</u> <u>Genetics Consortium (ADGC)</u>. Although this approach reduces the number of loci eligible for fine-mapping, the results are robust and reproducible. We obtained and reprocessed 38 myeloid epigenomic datasets to generate standardized annotations ^14, 32–36^, selected the ones that overlapped with active enhancers in myeloid cells and quantified their enrichment at active enhancers in each locus (Supplementary Figure 3). We then used PAINTOR with significantly enriched annotations (see Methods) to prioritize candidate causal variants and selected those with posterior probabilities of at least 10%. To investigate the biological mechanisms by which these variants could exhibit their effects, we screened for disruption or creation of binding motifs for TFs expressed in human microglia (TPM ≥1) ^14^ using motifbreakR^37^. We identified candidate non-coding functional variants in the BIN1, MS4A and ZYX loci and propose their likely mechanism of action (Supplementary Table 7). As an example, in the BIN1 locus we identified two independent AD GWAS signals. One of these signals is associated with a stimulation-dependent eQTL variant rs6733839-T that resides in a PU.1 binding site in microglia, disrupts the binding motif of the MEF2 transcription factor, likely acting as a binding partner for PU.1 at that site, and is an eQTL for BIN1 in monocytes stimulated with IFN-*γ*. The other variant (rs13025717-T) also resides in a PU.1 binding site, is an eQTL for BIN1 in all three myeloid cell types studied here and a binding QTL for PU.1 in a B-lymphoblastoid cell line (GM12878). This variant likely affects PU.1 binding by disrupting motifs of its binding partners, such as SP1 and KLF4 ^38, 39^. For the loci that were not significant in the ADGC GWAS (but were significant in the IGAP GWAS), we employed an alternative strategy for fine-mapping. Briefly, using a block partitioning algorithm ^40^, variant tagging algorithm ^41^ and conditional analyses ^42^ we were able to identify LD blocks and their tagging variants that independently contribute to the AD GWAS signal (see Methods). We overlapped these variants with active enhancer annotations and eQTL effects in monocytes (obtained from the Cardiogenics and Fairfax studies) and macrophages (obtained from the Cardiogenics study) to prioritize variants with regulatory potential in myeloid cells, then screened them for disruption or creation of binding motifs for TFs expressed in human microglia (TPM ≥1) ^14^ (see Methods). Using this approach, we identified candidate non-coding causal variants in the GPR141, RABEP1, SPI1, SPPL2A/AP4E1 and TP53INP1 loci and propose their likely mechanism of action (Supplementary Table 7). We performed conditional analyses using candidate functional variants as covariates and confirmed that they do indeed tag the entire AD GWAS signal in their respective loci (Supplementary Figure 4). SNP-targeted SMR analyses also confirmed that all candidate functional variants drive the causal association between gene expression levels in myeloid cells and AD risk in their respective loci (Supplementary Tables 8).

### A candidate causal variant in the MS4A locus disrupts an anchor CTCF binding site, likely altering chromatin looping and activity to increase MS4A6A gene expression and AD risk in myeloid cells and hiPSC-derived microglia

One of the prioritized candidate functional variants in the MS4A locus, the rs636317-T AD risk-increasing allele (11:60019150:C:T in GRCh37.p13 coordinates), resides in a CTCF binding site (Figure 4b (ii)). CTCF binding sites serve as anchors for long-range chromatin loops and this protein plays a pivotal role in determining the spatial organization of chromatin to regulate gene expression ^43^. The CTCF motif is highly evolutionarily conserved, and previous studies have shown that single point mutations in this motif can lead to dramatic dysregulation of chromatin looping and activity ^43^. We further confirmed that rs636317-T not only resides in a CTCF ChIP-Seq peak in monocytes, but also breaks the CTCF binding consensus sequence (Figure 4b (iii) and is a binding QTL for CTCF in a B-lymphoblastoid cell line (GM12878). Additionally, the CTCF binding QTL signal in GM12878 ^44^ has a 97.9 % probability of colocalization with AD risk alleles at this locus. Given that rs636317-T is predicted to disrupt a CTCF binding site, we hypothesized that this SNP destroys one of the two anchor CTCF binding sites in a chromatin loop, leading to altered chromatin architecture and activity in the locus, which in turn leads to upregulation of MS4A6A expression and increased AD risk. rs636317-T is an hQTL for multiple active enhancers in monocytes and a strong eQTL for MS4A6A in monocytes and macrophages, reinforcing the hypothesis that rs636317-T causes epigenetic dysregulation in the locus, which in turn leads to increased expression of MS4A6A (Figure 4d). Examination of promoter-capture Hi-C interactions in this region in monocytes and macrophages identified a conserved chromatin loop that connects the MS4A6A promoter to an inactive enhancer approximately 360 kilobases away (Figure 4a (vi)). Importantly, examination of ChIA-PET interactions for CTCF and RAD21 (a component of the cohesin complex often colocalized with CTCF at anchor sites to form chromatin loops ^43^) in GM12878 identified a chromatin loop that connects a CTCF/RAD21 anchor site in the same inactive enhancer to the CTCF/RAD21 anchor site likely disrupted by rs636317-T (Figure 4a (vii-ix)). This arrangement suggests that rs636317-T may alter chromatin architecture in such a way that the promoter of MS4A6A may lose its interaction with the inactive enhancer mentioned above and instead fall under the influence of other regulatory elements that may boost MS4A6A expression in myeloid cells. Another established role of CTCF is the separation of regions of inner condensed chromatin and outer open chromatin domains, marking repressed and active regions, respectively ^43^.

**Figure 4.**
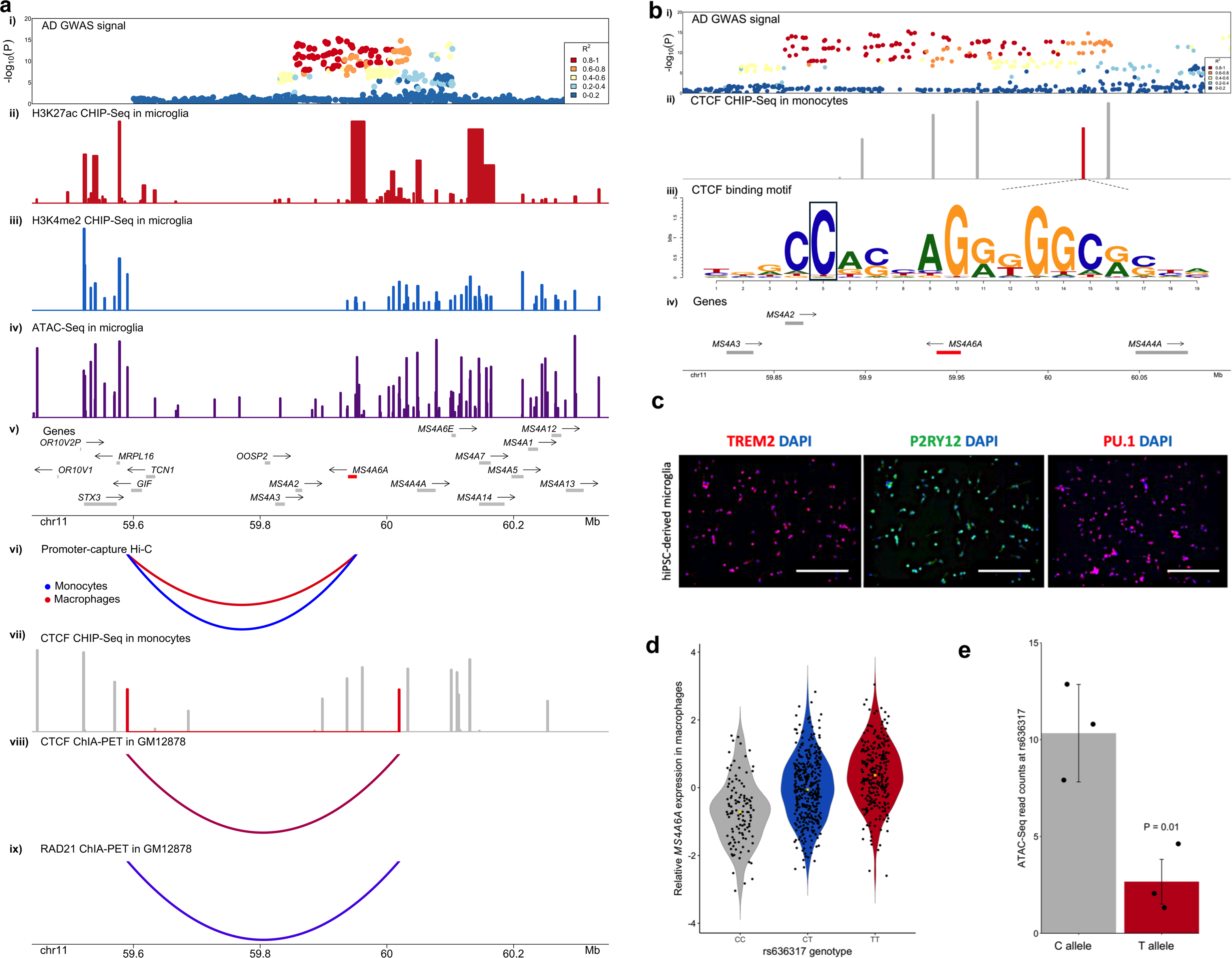
A candidate causal variant in the MS4A locus disrupts an anchor CTCF binding site, likely altering chromatin looping and activity to increase MS4A6A expression and AD risk in myeloid cells and hiPSC-derived microglia. a. i) AD GWAS signal in the MS4A locus. ii) H3K27ac peaks in microglia. iii) H3K4me2 peaks in microglia. iv) ATAC-Seq peaks in microglia. v) Genes that reside in the locus are plotted. Putative AD risk genes are highlighted in red. The arrow indicates the direction of transcription, while the bar indicates the gene body. vi) Promoter-capture Hi-C interactions between the MS4A6A promoter and a distal inactive enhancer in monocytes (blue) and macrophages (red). vii) CTCF ChIP-Seq peaks in monocytes. The peaks highlighted in red are anchor CTCF binding sites for the chromatin loop. viii) CTCF ChIA-PET interactions in GM12878. ix) RAD21 ChiA-PET interaction in GM12878. b. i) AD GWAS signal in the MS4A locus. ii) CTCF ChIP-Seq peaks in monocytes. The peak highlighted in red is an anchor CTCF binding site for a chromatin loop and contains the candidate causal variant (rs636317-T). iii) A CTCF binding motif resides in the CTCF ChIP peak highlighted in red in ii). The candidate causal variant (rs636317-T) resides in position 5 (boxed) of this motif and is predicted to disrupt CTCF binding. iv) Genes that reside in the locus are plotted. Putative AD risk genes are highlighted in red. The arrow indicates the direction of transcription, while the bar indicates the gene body. c. Representative immunofluorescent images of microglial markers (TREM2, P2RY12 and PU.1) confirming differentiation of hiPSC-derived microglia. Scale bar = 300μm. d. Relative expression of MS4A6A in macrophages increases in a rs636317-T allele dose-dependent manner. Each dot represents relative expression level of MS4A6A in each individual, while the yellow dot represents the median. e. Allelic imbalance of chromatin accessibility at the rs636317 site is observed in hiPSC-derived microglia. Mean ATAC-Seq read counts are plotted for the protective (C) and risk-increasing (T) alleles; the dots represent each individual and error bars represent standard errors of the mean. The protective allele (C) shows significantly more ATAC-Seq read counts than the risk-increasing allele (T) (P-value=0.01, one-sided t-test), which is consistent with the hypothesis that the presence of the rs636317 AD risk-increasing allele leads to disruption of CTCF binding.

Hence, we examined the density of epigenetic signals within and outside the Hi-C loop boundaries in microglia and observed that chromatin activity within the loop is repressed (Figure 4a (ii-iv)). To gather additional experimental evidence in support of our hypothesis, we investigated whether the C to T variation at rs636317 results in differential chromatin accessibility at this site in human microglia. To accomplish this, we generated hiPSC-derived microglia (Figure 4c) from 3 subjects heterozygous at rs636317, performed ATAC-Seq and quantified the number of reads that correspond to the protective and risk-increasing alleles. We observed a significant difference in the number of ATAC-Seq reads overlapping rs636317 with the protective allele (C) compared to the risk-increasing allele (T) (P-value=0.01, one-sided t-test) (Figure 4e), reinforcing the hypothesis that presence of the rs636317 AD risk-increasing allele leads to disruption of CTCF binding, decreased chromatin accessibility at this site, altered chromatin looping in the locus, and increased expression of MS4A6A in microglia.

### Discussion

In this study we report, for the first time, an integration of AD GWAS data with epigenomic and transcriptomic datasets from myeloid cells to nominate candidate causal variants, regulatory elements, genes and pathways and thus inform a mechanistic understanding of AD genetics and pathobiology for the formulation of novel therapeutic hypotheses. Previous studies have shown that myeloid cells are the most disease-relevant cell type for AD ^7, 12^ and our own earlier study showed an enrichment of AD SNP heritability in myeloid-specific epigenomic annotations including the PU.1 cistrome ^11^. Here we have extended these observations to demonstrate that AD risk alleles are specifically enriched in active enhancers of monocytes, monocyte-derived macrophages and microglia. Concordant with previous studies ^14, 19^, we show that PU.1, C/EBP, CTCF and RUNX binding motifs are overrepresented in open chromatin regions associated with active enhancers in all three myeloid cell types, while MEF2 transcription factor binding motifs are specifically overrepresented in open chromatin regions associated with microglial active enhancers. To identify transcription factor binding sites burdened by AD risk variants, we stratified open chromatin regions that overlapped with myeloid active enhancers by the presence of cognate consensus motifs for the TFs mentioned above and quantified the enrichment of AD risk alleles in these subsets. A significant enrichment was observed in PU.1 binding sites in all three myeloid cell types, while MAF binding sites were specifically enriched in macrophages and microglia. Furthermore, a significant enrichment of AD risk alleles was observed in SMAD, USF and SP1 binding sites in microglia. These results strongly suggest that AD risk variants are likely to modify disease susceptibility, at least in part, by modulating the binding of TFs to their cognate sequences in myeloid enhancers thus affecting their activity and in turn leading to causal target gene expression dysregulation. Although the global enrichment of AD risk alleles in active enhancers of myeloid cells narrows the search space for causal regulatory elements, identifying the target genes of these enhancers would directly point to candidate causal genes in AD risk loci.

In this study we used two complementary approaches to prioritize candidate causal target genes of myeloid active enhancers in AD risk loci. First, we mapped AD risk enhancers to their target genes in myeloid cells using chromatin interactions (Hi-C) and eQTL datasets from monocytes and macrophages. Using this approach, we identified previously nominated AD risk genes (BIN1 ^24^, MS4A6A ^11^, SPI1 ^11^) as well as novel candidate causal genes including AP4E1, APPB3, RIN3, TP53INP1 and ZYX in fourteen loci. In a subset of AD risk loci we report shared active enhancers that interact with multiple target gene promoters to regulate their expression. This could reflect either multiple causal genes within the locus or a single causal gene and several risk neutral genes that show association by due to transcriptional co-regulation. Additional evidence will be necessary to distinguish between these two possibilities and prioritize one or more genes at these loci. Second, we used SMR to infer the causal relationships between chromatin activity at myeloid enhancers with target gene expression regulation and AD risk modification. We sequentially studied the causal path linking enhancer activity with gene expression in myeloid cells using myeloid hQTLs as the exposure and myeloid eQTLs as the outcome, followed by myeloid eQTLs as the exposure and AD diagnosis as the outcome to identify active enhancers that likely modulate AD risk by regulating the expression of causal genes in myeloid cells. Using this approach, we identified previously nominated AD risk genes MS4A4A ^11^, MS4A6A ^11^, SPI1 ^11^, as well as novel candidate causal genes AP4E1, AP4M1, PILRA, RABEP1, SPPL2A, TP53INP1, ZKSCAN1, and ZYX in ten loci. Importantly, these two analytical approaches yielded largely overlapping results and led to the nomination of several candidate causal genes in sixteen loci (Figure 5). Moreover, in some of these loci the AD risk enhancers interact with the promoter of the same genes that show a statistically significant causal association through SMR (i.e., BIN1, SPI1, TP53INP1 and ZYX), reinforcing their regulatory potential, target gene nomination and disease relevance.

**Figure 5.**
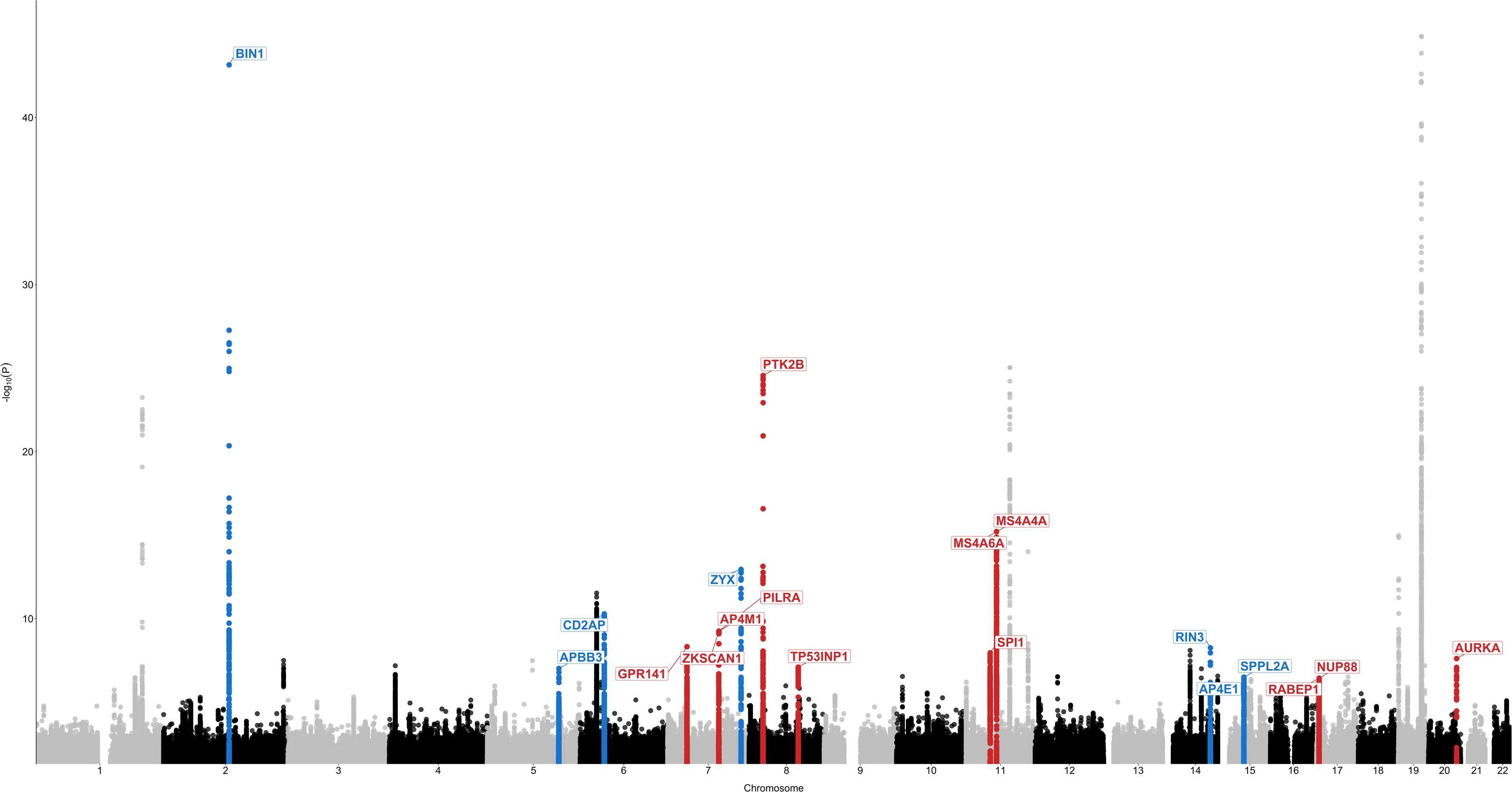
Candidate causal genes nominated through both Hi-C and SMR approaches in sixteen loci. Manhattan plot depicts the IGAP GWAS signal with putative AD risk genes assigned to each locus through both Hi-C and SMR approaches. Red indicates that increased expression of the gene is predicted to increase risk for AD. Blue indicates that decreased expression of the gene is predicted to increase risk for AD. The TREM2 and ABCA7 loci are not shown since TREM2 and ABCA7 are well established AD risk genes in their respective loci due to well replicated associations of AD risk with rare loss-of-function mutations in these genes^3, 6, 64^.

Remarkably, many of the novel candidate causal genes that we identified in this study are functionally related to the endolysosomal system. For example, ZYX encodes a zinc-binding phosphoprotein that localizes to early endosomes and phagosomes in IFN-*γ*-activated macrophages ^45^ and drives their intracellular movement by assembling actin filament rocket tails ^46^. RIN3 (Ras And Rab Interactor 3) encodes a member of the RIN family of RAS and RAB effectors that interacts and localizes with BIN1 to early endosomes ^47^. Like other RIN family members, RIN3 has guanine nucleotide exchange factory (GEF) activity for RAB5 GTPases ^47^, which are required for early endosome and phagosome biogenesis and function. Interestingly, RABEP1 (Rab-GTPase binding effector protein 1) also encodes a RAB5 effector protein that is required for early endosome membrane fusion and trafficking ^48^. Two other novel candidate AD risk genes that we nominated in this study, AP4E1 and AP4M1, encode two of the four subunits of the heterotetrameric adaptor protein complex 4 (AP-4), which is required for the sorting of transmembrane proteins like APP from the trans-Golgi network (TGN) to endosomes^49^. Interestingly, APBB3 has also been shown to bind to the intracellular domain of APP and is thought to play a role in the internalization of APP from the cell surface into endosomes where it is cleaved by membrane-embedded aspartyl proteases BACE1 and ɣ-secretase to generate the amyloid β peptide ^50, 51^. Another novel candidate AD risk gene that we nominate in this study, SPPL2A, encodes a transmembrane aspartyl protease that localizes to late endosomes and lysosomes and cleaves substrates involved in immunity and neurodegeneration ^52–54^. Finally, TP53INP1 regulates the stability and transcriptional activity of p53, and has been implicated in the phagocytic clearance of apoptotic cells (efferocytosis) ^55, 56^, a hallmark function of macrophages for the maintenance of tissue homeostasis and the resolution of inflammation. All of these genes are highly or selectively expressed in microglia in the brain ^14^. Taken together, our findings implicate dysfunction of the endolysosomal system in myeloid cells (as opposed to neurons ^57^) in the etiology of AD. Previous human genetic findings reinforce our conclusion. For example, a rare variant in the 3′ UTR of RAB10, a member of the RAB family of small GTPases that are critical regulators of membrane trafficking and vesicular transport, confers resilience to AD ^58^. Furthermore, coding variants that increase risk for AD have been identified in SORL1^4, 59^, a member of the vacuolar protein sorting 10 (VPS10)- domain-containing receptor family and the low density lipoprotein receptor (LDLR) family of APOE receptors that is expressed primarily in microglia in the brain ^14^ and plays important roles in the endolysosomal system and APP processing ^57^.

To fine-map the AD risk enhancers identified in this study and thus nominate candidate causal variants, we conducted Bayesian fine-mapping in the three loci that were significantly associated with AD risk in the ADGC GWAS (BIN1, MS4A and ZYX), followed by functional in silico screening of the candidate causal variants for disruption/creation of TF binding motifs. We also fine-mapped the loci that did not reach significance in the ADGC GWAS (but were significant in the IGAP GWAS) and identified candidate causal variants in the GPR141, RABEP1, SPI1, SPPL2A/AP4E1 and TP53INP1 loci. Taken together, we have identified putative functional variants that tag the entire AD GWAS signals at these loci, and likely affect disease risk by altering the DNA binding motifs of transcription factors that modulate the activity of enhancers which in turn regulate the expression of causal genes to ultimately steer myeloid cells like microglia toward neurotoxic and/or away from neuroprotective phenotypes. Finally, we experimentally validated one of these candidate functional variants in the MS4A locus, which disrupts CTCF binding to one of two anchor sites of a repressive chromatin loop, leading to increased MS4A6A expression and AD risk.

In summary, this study reveals a link between enhancer activity, gene expression and AD risk in monocytes, macrophages and microglia, proposes the molecular mechanism of action of candidate functional variants in several AD risk loci, identifies specific AD risk enhancers that are burdened by these variants and regulate causal gene expression, which in turn most likely modulates disease susceptibility by altering the biology of myeloid cells. We highlight the coalescence of candidate causal genes in the endolysosomal system of myeloid cells and underscore its importance in the etiology of AD.

## Online Methods

### Processing of ChIP-Seq and ATAC-Seq data and peak calling

Relevant ChIP-Seq studies were found through Gene Expression Omnibus (GEO). Fastq files were obtained from Sequence Read Archive (SRA) and FASTQC was used for quality control of the files. Poor quality samples were discarded. Technical replicates were merged and the files were trimmed with trimgalore (see URLs). Bowtie2 ^60^ was used for alignment for both single and paired-end files and resulting sam files were filtered by MAPQ score. Samtools ^61^ were used to remove PCR duplicates and MACS2^62^ was used to call peaks. ATAC-Seq peaks were called using the following command: “callpeak -t file.sam -f SAM --nomodel --shift -37 --extsize 73 -g hs -q 0.01 -n filename --outdir output_dir/”. PU.1 ChIP-Seq peaks were called using the following command: callpeak -t case.sam -c input.sam -f SAM -g hs -q 0.01 -n filename --outdir output_dir/”. Histone modifications ChIP-Seq peaks were called using the following command: “callpeak -t case.sam -c input.sam -f SAM --broad --broad-cutoff 0.01 -g hs -q 0.01 -n filename --outdir output_dir/”. Samtools mpileup function was used to quantify the number of reads that align to each allele.

### Stratification into promoter and enhancer regions

To identify optimal distance from TSS we used ChromHMM model of CD14+ monocytes from Roadmap Epigenomics project (see URLs) to visualize the distribution of active and primed promoters around the TSS. We observed a bimodal distribution around the TSS and found that -500 base pairs to 1000 base pairs window captures more than 60% of active promoters. To annotate the peaks with distance from TSS we used HOMER. We then split the H3K4me1/2 peaks into distal (further than 500 base pairs to the left OR further than 1000 base pairs to the right from the TSS) and proximal (between 500 base pairs to the left and 1000 base pairs to the right). We then used bedmap to filter H3K4me1/2 peaks by the presence of H3K27ac peak such that proximal H3K4me1/2 peaks with H3K27ac are active promoters, distal H3K4me1/2 peaks with H3K27ac are active enhancers, proximal H3K4me1/2 peaks without H3K27ac are primed promoters and distal H3K4me1/2 peaks without H3K27ac are primed enhancers.

### Partitioned SNP-heritability analysis

We used LD Score regression to estimate AD SNP heritability partitioned by epigenomic annotations using GWAS summary statistics (excluding the APOE (chr19:45000000– 45800000) and MHC/HLA (chr6:28477797–33448354) regions) in myeloid cells as described in the companion website (see URLs), while controlling for the 53 functional annotation categories of the full baseline model. GWAS summary statistics for AD ^16^ and Schizophrenia ^17^ (SCZ) were downloaded from the IGAP Consortium and Psychiatric Genomics Consortium websites respectively (see URLs). All epigenomic annotations were downloaded from SRA and preprocessed and the peaks were called as described in “Processing of ChIP-Seq data and peak calling”.

### De novo motif discovery

We used HOMER to perform de novo motif discovery in ATAC-Seq regions that reside in active enhancers in monocytes, macrophages and microglia. The following command was used to identify enriched motif sequences in these regions: findMotifsGenome.pl Peaks.bed hg19. -size given.

### Causal association analysis

We used SMR to infer causal associations between IGAP GWAS and QTL datasets^13^. We converted the summary statistics for monocyte H3K4me1 hQTLs obtained from BLUEPRINT epigenome project website (see URLs) and monocyte eQTLs from the Cardiogenics and Fairfax studies into BESD format (epi/esi/besd) as described in the SMR manual (see URLs). Allele frequencies and LD were estimated from the ADGC GWAS cohort individual-level genotype data. To conduct standard SMR analysis, we ran the following command: “smr --bfile reference_file --beqtl-summary Histone_besd_file_prefix --beqtl-summary eQTL_besd_file_prefix --out output_prefix”. The results were filtered for FDR of 10% using R. To conduct SNP-targeted SMR analysis, we ran the following command: “smr --bfile reference_file --gwas-summary gwas_summary_file --beqtl-summary eQTL_besd_fie_prefix --target-snp rs12345 --out output_prefix”.

### Colocalization analysis

We used coloc (coloc.abf function) to perform colocalization analyses between IGAP GWAS and QTL datasets^27^.

### Conditional and haplotype analyses

We used GCTA-COJO^42^to conduct multi-SNP based conditional analyses using IGAP GWAS summary statistics data and ADGC GWAS cohort individual-level genotype data as a reference panel (see URLs). Allele frequencies and LD were estimated from the ADGC GWAS cohort individual-level genotype data. To conduct the conditional analysis we ran the following command: “gcta64 --bfile reference_file --maf 0.05 --cojo-file IGAP_GWAS_summary_statistics--cojo-cond list_of_snps --out output_prefix”. To construct haplotype blocks and examine SNP clustering, we used BigLD ^40^ which is provided as an R package (see URLs). We prepared the genotype file, which contained genotypes of individuals for each SNP, and the SNP information file that contained chromosome, position, reference and alternative allele information for each SNP. We then used CLQ algorithm provided within BigLD package for SNP clustering and Big_LD for haplotype block construction. We used LDblockHeatmap function to visualize the blocks identified by BigLD along with SNP clusters.

### Fine-mapping analysis

We used PAINTOR to conduct fine-mapping of AD risk loci. PAINTOR is a Bayesian fine-mapping method that leverages functional annotations through an Empirical Bayes prior ^30^. The input files for PAINTOR_v3.1 were prepared as described on the PAINTOR website and ADGC GWAS summary statistics along with individual-level genotype data were used for fine-mapping (see URLs). The reprocessed epigenomic annotations were all used to quantify enrichment at each locus. To quantify the annotation enrichments the following command was used: “python AnnotateLocus.py --input list_of_annotation_directories --locus locus_prefix --out output_prefix --chr chr --pos pos”. To classify the annotations as enriched or not, we computed the relative probability for a SNP to be causal given that it resides in the annotation as follows: Baseline prior probability of a SNP to be causal = 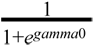, where gamma0 is the effect size estimate for the baseline without the annotation. Prior probability of a SNP to be causal given it is in the annotation =, 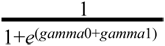, where gamma1 is the effect size estimate for the annotation. From the formula above it is evident that it is desirable that the significant annotation has a negative effect size estimate. We, thus, compute the relative probability of a SNP to be causal given that it is in the annotation in the following manner: The relative probability of a SNP to be causal given that it is in the annotation = 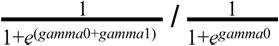. We deemed the annotation to be significant if the relative probability of a SNP to be causal given that it is in the annotation was greater than 1.

### TF binding motif disruption/creation analysis

We used motifbreakR to predict the impact of AD risk variants on transcription factor binding^37^. We used HOCOMOCO to screen for TFBMs and a P-value significance threshold of 5×10^-5^ as advised by the authors of the package.

### Prioritization of candidate causal variants in loci that are not significant in ADGC GWAS

For each locus, we first constructed LD blocks using BigLD package ^40^. We also constructed haplotypes using Haploview to assess consistency of haplotype blocks^63^. We then used the tagger functionality within Haploview to identify tagging variants for these blocks. We conducted conditional analyses by adding each of the tagging variants sequentially to the model to identify independent LD blocks and a set of tagging variants that account for the entire GWAS signal at each locus. We conducted a motif disruption/creation analysis on the variants within the disease-associated blocks and selected the variants that are predicted to strongly disrupt or create binding sites of transcription factors that are expressed in myeloid cells (TPM≥1)^14^. We further overlapped the variants within the blocks with our active enhancer annotations in monocytes, macrophages and microglia. We then screened the remaining variants for eQTLs in monocytes and macrophages from the Cardiogenics and Fairfax studies. Once candidate causal variants were selected, we conducted conditional analyses to make sure that they do indeed tag the entirety of the GWAS signal in the locus.

### Generation of hiPSC microglia for ATAC-Seq analysis

hiPSC-derived microglia were generated from patient lines following the protocol as described (Abud et al., 2017). For the ATAC-Seq analysis, hiPSC-derived microglia (50K cells) from each patient line were collected and processed as described (Buenrostro, 2014). Samples were either processed at New York Genome Center or at UCI’s Genomics High-Throughput Facility and sequenced as 50 bp paired-end reads on a HiSeq 2500 and 100 bp paired-end reads on a HiSeq 4000, respectively. The consent for reprogramming patient somatic cells to hiPSC was carried out on protocol 2013-9561 (UCI), laboratory protocol 2017-1061 (UCI) and protocol ESCRO 19-04 (Mount Sinai).

### Immunocytochemistry

Cells were fixed with 4% paraformaldehyde in PBS at 4°C for 10 min. Cells were permeabilized with 1.0% Triton in PBS at room temperature for 15 min and blocked in 5% donkey serum with 0.1% Triton in PBS at room temperature for 30 min. Primary antibodies were used at 10 µg/mL anti-TREM2 (R&D, AF1828), 1:1,000 anti-P2RY12 (Sigma, HPA014518), and 1:100 anti-PU.1 (Cell Signaling, 2266). Secondary antibodies were used at 1:300 Alexa donkey 488 and 568 anti-rabbit, mouse, or chicken (Life Technologies). DAPI (4’,6-diamidino-2-phenylindole, 0.5 μg/mL) was used to visualize nuclei. Images were acquired using a Leica Fluorescence Microscope.

### Data availability

All sequencing files and processed peaks for hiPSC-derived microglia ATAC-Seq will be deposited to the Gene Expression Omnibus once the manuscript is accepted for publication. The following studies obtained from GEO were used for the analyses presented in this paper: GSE29611, GSE85245, GSE100380, GSE66594, GSE85245. DbGAP accession study number for the human microglia dataset is phs001373.v1.p1. The genotype and phenotype data from ADGC are available under phs000372.v1.p1 dbGAP study accession number.

### Code availability

Although we have used the software cited in this manuscript with default parameters or minor changes, code for these analyses is available upon request.

## Acknowledgments

We thank the Cardiogenics (European Project reference LSHM-CT-2006-037593) project for providing summary statistics data for eQTL analyses in monocytes and monocyte-derived macrophages. We thank the New York Genome Center for sequencing the hiPSC-derived microglia samples.

## Author contributions

A.M.G., E.M. and G.N. conceived and designed the experiments. G.N, M.K, H.C, K.H and A.G.E performed data analyses. J.TCW. and E.M.A carried out the hiPSC-derived microglia studies under the supervision of W.W.P. P.R., J.F. and J. X. performed ATACseq expts. E.M and A.M.G. supervised data analysis. G.N, E.M. and A.M.G. wrote and edited the manuscript. All authors read and approved the manuscript.

## Competing interests

AMG has consulted for Eisai, Biogen, Pfizer, AbbVie, Cognition Therapeutics and GSK, she also served on the SAB at Denali Therapeutics from 2015-2018. This work was funded by grants from the NIH: U01AG052411 (AMG), RF1AG054011 (AMG), U01AG058635 (AMG), NIA K01AG062683 (J.TCW.), AG016573 (WWP), F31 AG059337-01 (A.G.E), The JPB Foundation, The Robert and Renee Belfer Foundation.

## Notes

#### Summary of Updates

Figure 4 revised

## References

1. Dementia statistics | Alzheimer’s Disease International. Available at: https://www.alz.co.uk/research/statistics. (Accessed: 7th April 2019)

2. Efthymiou, A. G. & Goate, A. M. Late onset Alzheimer’s disease genetics implicates microglial pathways in disease risk. Mol. Neurodegener. 12, 43 (2017).

3. Jonsson, T. et al. Variant of TREM2 associated with the risk of Alzheimer’s disease. N. Engl. J. Med. 368, 107–116 (2013).

4. Vardarajan, B. N. et al. Coding mutations in SORL1 and Alzheimer disease. Ann. Neurol. 77, 215–227 (2015).

5. Sims, R. et al. Rare coding variants in PLCG2, ABI3, and TREM2 implicate microglial-mediated innate immunity in Alzheimer’s disease. Nat. Genet. 49, 1373–1384 (2017).

6. Steinberg, S. et al. Loss-of-function variants in ABCA7 confer risk of Alzheimer’s disease. Nat. Genet. 47, 445–447 (2015).

7. Hansen, D. V., Hanson, J. E. & Sheng, M. Microglia in Alzheimer’s disease. The Journal of Cell Biology 217, 459–472 (2018).

8. Kunkle, B. W. et al. Meta-analysis of genetic association with diagnosed Alzheimer’s disease identifies novel risk loci and implicates Abeta, Tau, immunity and lipid processing. doi:10.1101/294629

9. Marioni, R. E. et al. GWAS on family history of Alzheimer’s disease. Transl. Psychiatry 8, 99 (2018).

10. Jansen, I. E. et al. Genome-wide meta-analysis identifies new loci and functional pathways influencing Alzheimer’s disease risk. Nat. Genet. 51, 404–413 (2019).

11. Huang, K.-L. et al. A common haplotype lowers PU.1 expression in myeloid cells and delays onset of Alzheimer’s disease. Nat. Neurosci. 20, 1052–1061 (2017).

12. Finucane, H. K. et al. Heritability enrichment of specifically expressed genes identifies disease-relevant tissues and cell types. Nat. Genet. 50, 621–629 (2018).

13. Zhu, Z. et al. Integration of summary data from GWAS and eQTL studies predicts complex trait gene targets. Nat. Genet. 48, 481–487 (2016).

14. Gosselin, D., et al. An environment-dependent transcriptional network specifies human microglia identity. 3222, (2017).

15. Finucane, H. K. et al. Partitioning heritability by functional annotation using genome-wide association summary statistics. Nat. Genet. 47, 1228 (2015).

16. Lambert, J.-C. Meta-Analysis of 74,046 Individuals Identifies 11 New Susceptibility Loci for Alzheimer’s Disease. Nat. Genet. 45, 1452–1458 (2013).

17. Cross-Disorder Group of the Psychiatric Genomics Consortium. Identification of risk loci with shared effects on five major psychiatric disorders: a genome-wide analysis. Lancet 381, 1371–1379 (2013).

18. Javierre, B. M. et al. Lineage-Specific Genome Architecture Links Enhancers and Non-coding Disease Variants to Target Gene Promoters. Cell 167, 1369–1384.e19 (2016).

19. Lavin, Y. et al. Tissue-resident macrophage enhancer landscapes are shaped by the local microenvironment. Cell 159, 1312–1326 (2014).

20. Gosselin, D. et al. Environment drives selection and function of enhancers controlling tissue-specific macrophage identities. Cell 159, 1327–1340 (2014).

21. Garnier, S. et al. Genome-Wide Haplotype Analysis of Cis Expression Quantitative Trait Loci in Monocytes. PLoS Genet. 9, e1003240 (2013).

22. Fairfax, B. P. et al. Innate immune activity conditions the effect of regulatory variants upon monocyte gene expression. Science (2014). doi:10.1126/science.1246949

23. Reitz, C. et al. Variants in the ATP-Binding Cassette Transporter (ABCA7), Apolipoprotein E ɛ4, and the Risk of Late-Onset Alzheimer Disease in African Americans. JAMA 309, 1483–1492 (2013).

24. Chapuis, J. et al. Increased expression of BIN1 mediates Alzheimer genetic risk by modulating tau pathology. Mol. Psychiatry 18, 1225–1234 (2013).

25. Raj, T. et al. Integrative transcriptome analyses of the aging brain implicate altered splicing in Alzheimer’s disease susceptibility. Nature Genetics 50, 1584–1592 (2018).

26. Chen, L. et al. Genetic Drivers of Epigenetic and Transcriptional Variation in Human Immune Cells. Cell 167, 1398–1414.e24 (2016).

27. Giambartolomei, C. et al. Bayesian test for colocalisation between pairs of genetic association studies using summary statistics. PLoS Genet. 10, e1004383 (2014).

28. Ward, L. D. & Kellis, M. HaploReg: a resource for exploring chromatin states, conservation, and regulatory motif alterations within sets of genetically linked variants. Nucleic Acids Res. 40, D930–4 (2012).

29. Rathore, N. et al. Paired Immunoglobulin-like Type 2 Receptor Alpha G78R variant alters ligand binding and confers protection to Alzheimer’s disease. PLoS Genet. 14, e1007427 (2018).

30. Kichaev, G. et al. Integrating functional data to prioritize causal variants in statistical fine-mapping studies. PLoS Genet. 10, e1004722 (2014).

31. Benner, C. et al. Prospects of Fine-Mapping Trait-Associated Genomic Regions by Using Summary Statistics from Genome-wide Association Studies. Am. J. Hum. Genet. 101, 539–551 (2017).

32. ENCODE Project Consortium. An integrated encyclopedia of DNA elements in the human genome. Nature 489, 57–74 (2012).

33. Novakovic, B. et al. β-Glucan Reverses the Epigenetic State of LPS-Induced Immunological Tolerance. Cell 167, 1354–1368.e14 (2016).

34. Park, S. H. et al. Type I interferons and the cytokine TNF cooperatively reprogram the macrophage epigenome to promote inflammatory activation. Nat. Immunol. 18, 1104–1116 (2017).

35. Kang, K. et al. Interferon-γ Represses M2 Gene Expression in Human Macrophages by Disassembling Enhancers Bound by the Transcription Factor MAF. Immunity 47, 235–250.e4 (2017).

36. Schmidt, S. V. et al. The transcriptional regulator network of human inflammatory macrophages is defined by open chromatin. Cell Res. 26, 151–170 (2016).

37. Coetzee, S. G., Coetzee, G. A. & Hazelett, D. J. motifbreakR: an R/Bioconductor package for predicting variant effects at transcription factor binding sites. Bioinformatics 31, 3847–3849 (2015).

38. Feng, X. et al. Sp1/Sp3 and PU.1 Differentially Regulate β5Integrin Gene Expression in Macrophages and Osteoblasts. J. Biol. Chem. 275, 8331–8340 (2000).

39. Feinberg, M. W. et al. The Kruppel-like factor KLF4 is a critical regulator of monocyte differentiation. EMBO J. 26, 4138–4148 (2007).

40. Kim, S. A., Cho, C.-S., Kim, S.-R., Bull, S. B. & Yoo, Y. J. A new haplotype block detection method for dense genome sequencing data based on interval graph modeling of clusters of highly correlated SNPs. Bioinformatics 34, 388–397 (2018).

41. de Bakker, P. I. W. et al. Efficiency and power in genetic association studies. Nat. Genet. 37, 1217–1223 (2005).

42. Yang, J. et al. Conditional and joint multiple-SNP analysis of GWAS summary statistics identifies additional variants influencing complex traits. Nat. Genet. 44, 369–75, S1–3 (2012).

43. Tang, Z. et al. CTCF-Mediated Human 3D Genome Architecture Reveals Chromatin Topology for Transcription. Cell 163, 1611–1627 (2015).

44. Ding, Z. et al. Quantitative genetics of CTCF binding reveal local sequence effects and different modes of X-chromosome association. PLoS Genet. 10, e1004798 (2014).

45. Trost, M. et al. The phagosomal proteome in interferon-gamma-activated macrophages. Immunity 30, 143–154 (2009).

46. Southwick, F. S., Li, W., Zhang, F., Zeile, W. L. & Purich, D. L. Actin-based endosome and phagosome rocketing in macrophages: activation by the secretagogue antagonists lanthanum and zinc. Cell Motil. Cytoskeleton 54, 41–55 (2003).

47. Kajiho, H. et al. RIN3: a novel Rab5 GEF interacting with amphiphysin II involved in the early endocytic pathway. J. Cell Sci. 116, 4159–4168 (2003).

48. Stenmark, H., Vitale, G., Ullrich, O. & Zerial, M. Rabaptin-5 is a direct effector of the small GTPase Rab5 in endocytic membrane fusion. Cell 83, 423–432 (1995).

49. Burgos, P. V. et al. Sorting of the Alzheimer’s disease amyloid precursor protein mediated by the AP-4 complex. Dev. Cell 18, 425–436 (2010).

50. Duilio, A., Faraonio, R., Minopoli, G., Zambrano, N. & Russo, T. Fe65L2: a new member of the Fe65 protein family interacting with the intracellular domain of the Alzheimer’s beta-amyloid precursor protein. Biochem. J 330 (Pt 1), 513–519 (1998).

51. Tanahashi, H. & Tabira, T. Molecular cloning of human Fe65L2 and its interaction with the Alzheimer’s beta-amyloid precursor protein. Neurosci. Lett. 261, 143–146 (1999).

52. Behnke, J. et al. Signal-peptide-peptidase-like 2a (SPPL2a) is targeted to lysosomes/late endosomes by a tyrosine motif in its C-terminal tail. FEBS Lett. 585, 2951–2957 (2011).

53. Schneppenheim, J. et al. The intramembrane protease SPPL2a promotes B cell development and controls endosomal traffic by cleavage of the invariant chain. J. Exp. Med. 210, 41–58 (2013).

54. Brady, O. A., Zhou, X. & Hu, F. Regulated intramembrane proteolysis of the frontotemporal lobar degeneration risk factor, TMEM106B, by signal peptide peptidase-like 2a (SPPL2a). J. Biol. Chem. 289, 19670–19680 (2014).

55. Shahbazi, J., Lock, R. & Liu, T. Tumor Protein 53-Induced Nuclear Protein 1 Enhances p53 Function and Represses Tumorigenesis. Front. Genet. 4, 80 (2013).

56. Yoon, K. W. et al. Control of signaling-mediated clearance of apoptotic cells by the tumor suppressor p53. Science 349, 1261669 (2015).

57. Small, S. A., Simoes-Spassov, S., Mayeux, R. & Petsko, G. A. Endosomal Traffic Jams Represent a Pathogenic Hub and Therapeutic Target in Alzheimer’s Disease. Trends Neurosci. 40, 592–602 (2017).

58. Ridge, P. G. et al. Linkage, whole genome sequence, and biological data implicate variants in RAB10 in Alzheimer’s disease resilience. Genome Medicine 9, (2017).

59. Raghavan, N. S. et al. Whole-exome sequencing in 20,197 persons for rare variants in Alzheimer’s disease. Ann Clin Transl Neurol 5, 832–842 (2018).

60. Langmead, B. & Salzberg, S. L. Fast gapped-read alignment with Bowtie 2. Nat. Methods 9, 357–359 (2012).

61. Li, H. et al. The Sequence Alignment/Map format and SAMtools. Bioinformatics 25, 2078–2079 (2009).

62. Zhang, Y. et al. Model-based analysis of ChIP-Seq (MACS). Genome Biol. 9, R137 (2008).

63. Barrett, J. C., Fry, B., Maller, J. & Daly, M. J. Haploview: analysis and visualization of LD and haplotype maps. Bioinformatics 21, 263–265 (2005).

64. Guerreiro, R. et al. TREM2 variants in Alzheimer’s disease. N. Engl. J. Med. 368, 117–127 (2013).

