## Supplementary figures and images for "Integration of Alzheimer’s disease genetics and myeloid genomics reveals novel disease risk mechanisms"

### Supplementary figures

#### Supplementary Figure 1

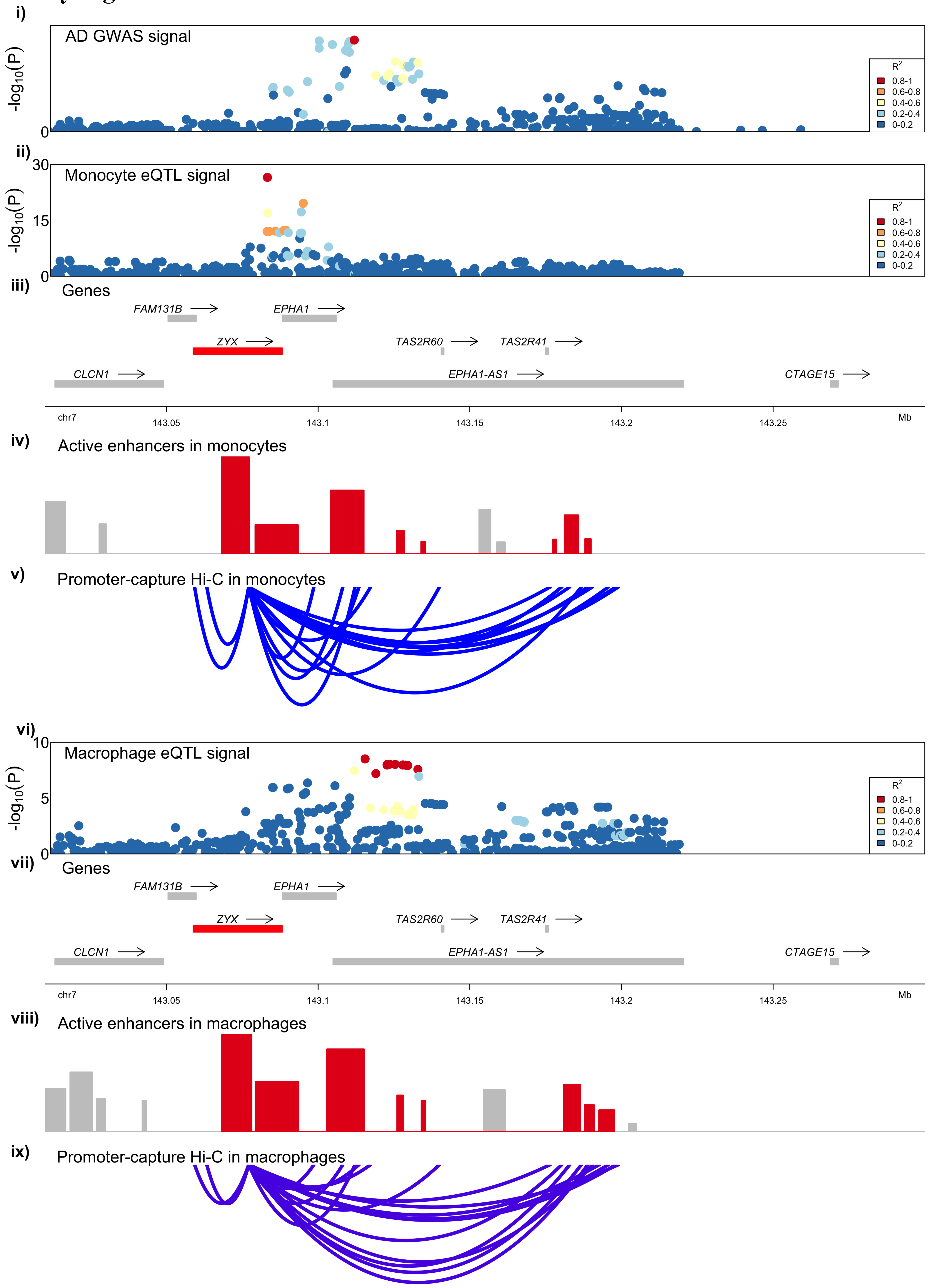

##### Supplementary Figure 2

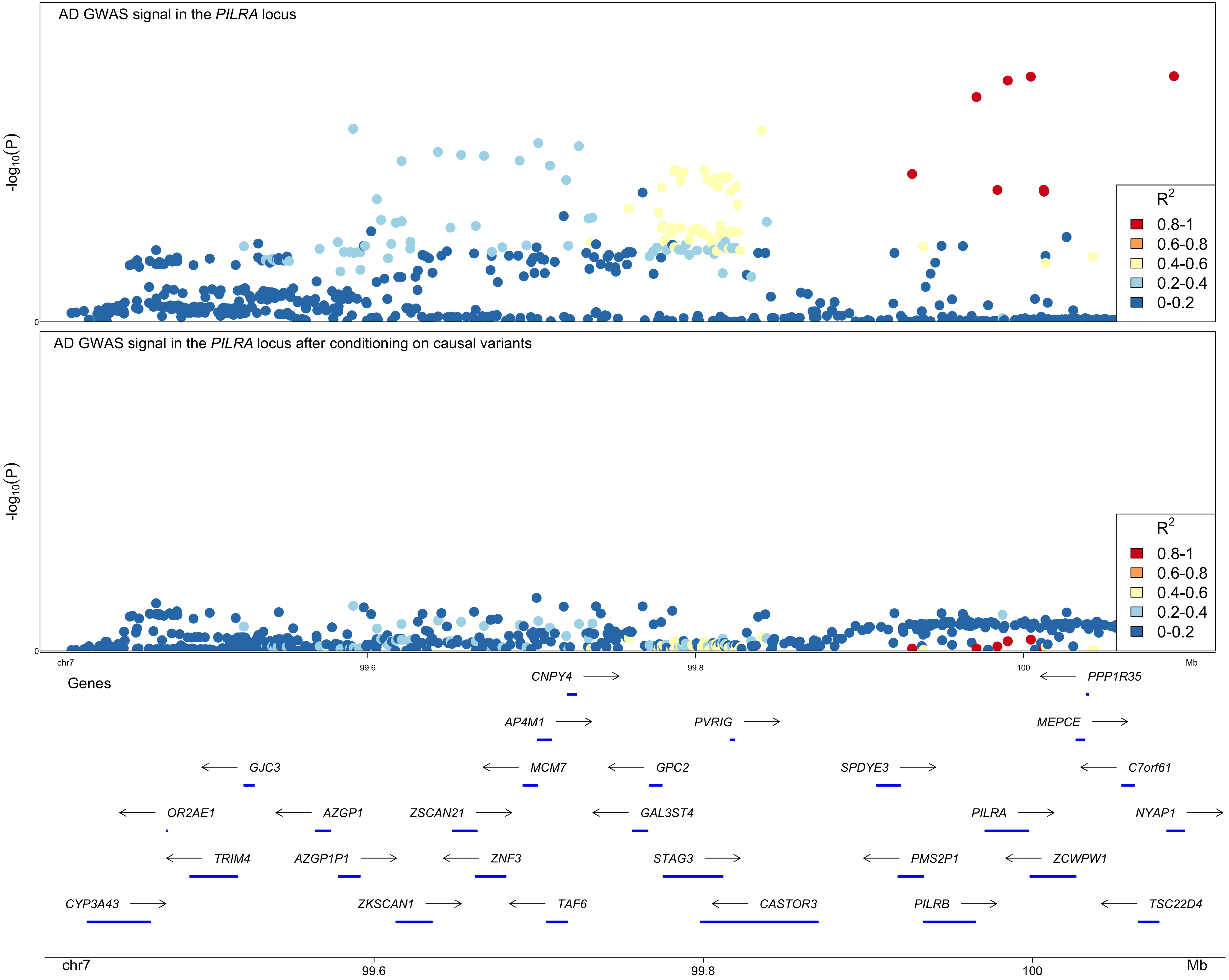

#### Supplementary Figure 3

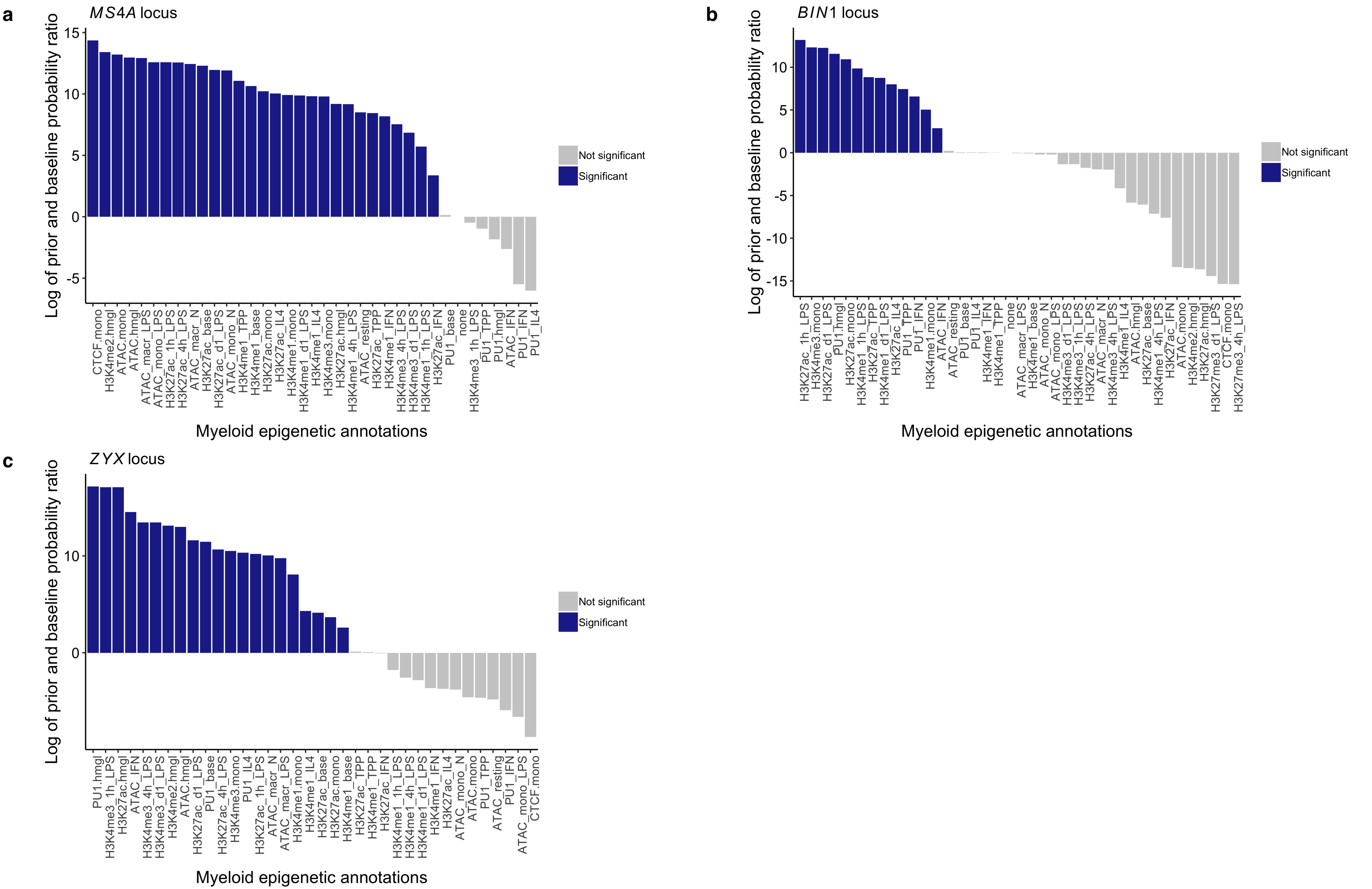

#### Supplementary Figure 4

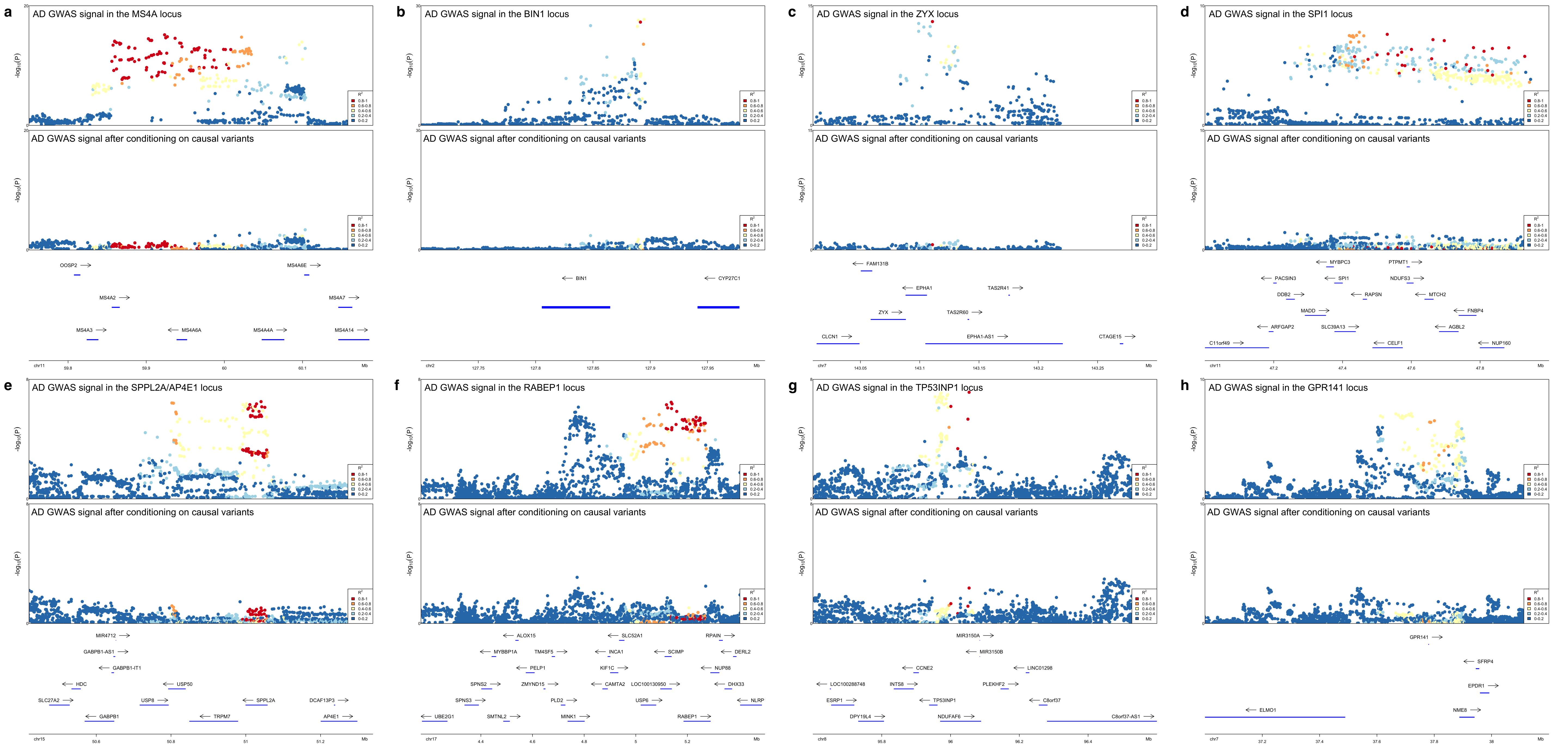
