## Supplementary figure and table legends for "Integration of Alzheimer’s disease genetics and myeloid genomics reveals novel disease risk mechanisms"

**Supplementary Figure Legends**

**Supplementary Figure 1.** i) AD GWAS signal in the *ZYX* locus. ii) eQTL signal for *ZYX* in monocytes obtained from the Cardiogenics study. iii) Genes that reside in the locus are plotted. Putative AD risk genes are highlighted in red. iv) Active enhancers in monocytes are plotted. Putative AD risk enhancers are highlighted in red. v) Promoter-capture Hi-C interactions between the *ZYX* promoter and AD risk enhancers in monocytes. vi) eQTL signal for *ZYX* in macrophages obtained from the Cardiogenics study. vii) Genes that reside in the locus are plotted. Putative AD risk genes are highlighted in red. viii) Active enhancers in macrophages are plotted. Putative AD risk enhancers are highlighted in red. ix) Promoter-capture Hi-C interactions between the *ZYX* promoter and AD risk enhancers in macrophages.

**Supplementary Figure 2.** Conditional analysis plots of the missense variant (rs1859788-A) in the *PILRA* locus.

**Supplementary Figure 3.** Log10 of prior and baseline probabilities ratio obtained from PAINTOR fine-mapping analysis of the a) *MS4A* b) *BIN1* and c) *ZYX* loci plotted for each myeloid epigenomic annotation tested. Annotations that are significantly enriched in the locus are colored in blue, while non-significant annotations are colored in grey (see Methods).

**Supplementary Figure 4.** Conditional analysis plots of the candidate causal variants listed in Supplementary Table 8 in the a) *MS4A*, b) *BIN1*, c) *ZYX*, d) *SPI1*, e) *SPPL2A/AP4E1*, f) *RABEP1*, g) *TP53INP1* and h) *GPR141* loci.

**Supplementary Table Legends**

**Supplementary Table 1.** HOMER *de novo* motif discovery results for ATAC-Seq regions that reside in active enhancers in monocytes (Sheet 1), macrophages (Sheet 2) and microglia (Sheet 3).

**Supplementary Table 2.** Colocalization analysis results for active enhancers in monocytes that contain AD risk alleles and hQTLs. PP.H3 = posterior probability for the hypothesis of independent AD GWAS and hQTL signals. PP.H4 = posterior probability for the hypothesis of colocalized AD GWAS and hQTL signals.

**Supplementary Table 3.** SMR analysis results for chromatin activity (hQTLs) and gene expression (eQTLs) in monocytes (obtained from the Cardiogenics study) at enhancers selected using coloc.

**Supplementary Table 4.** SMR analysis results for gene expression (eQTL) in monocytes (obtained from the Cardiogenics study) and AD risk at enhancers selected using coloc and SMR analysis as described for Supplementary Table 3.

**Supplementary Table 5.** SMR analysis results for chromatin activity (hQTLs) and gene expression (eQTLs) in monocytes (obtained from the Fairfax study) at enhancers selected using coloc.

**Supplementary Table 6.** SMR analysis results for gene expression (eQTL) in monocytes (obtained from the Fairfax study) and AD risk at enhancers selected using coloc and SMR analysis as described for Supplementary Table 5.

**Supplementary Table 7.** Fine-mapping analysis results in eight AD risk loci including the number of independent causal variants, the candidate causal variants, the proposed mechanisms of action, and the direction of the eQTL effects that increase AD risk.

**Supplementary Table 8.** SNP-targeted SMR analysis results for candidate causal variants listed in Supplementary Table 7.
